# Higher-order interactions among coinfecting parasites and a microbial mutualist impact disease progression

**DOI:** 10.1101/2021.03.25.437101

**Authors:** Kayleigh R. O’Keeffe, Anita Simha, Charles E. Mitchell

## Abstract

Interactions among parasites and other microbes within hosts can impact disease progression, yet study of such interactions has been mostly limited to pairwise combinations of microbes. Given the diversity of microbes within hosts, higher-order interactions among more than two microbial species may also impact disease. To test this hypothesis, we performed inoculation experiments that investigated interactions among two fungal parasites, *Rhizoctonia solani* and *Colletotrichum cereale*, and a systemic fungal endophyte, *Epichloë coenophiala*, within a grass host. Both pairwise and higher-order interactions impacted disease progression. While the endophyte did not directly influence *R. solani* growth or *C. cereale* symptom development, the endophyte modified the interaction between the two parasites. The magnitude of the facilitative effect of *C. cereale* on the growth of *R. solani* tended to be greater when the endophyte was present. Moreover, this interaction modification strongly affected leaf mortality. For plants lacking the endophyte, parasite co-inoculation did not increase leaf mortality compared to single-parasite inoculations. In contrast, for endophyte-infected plants, parasite co-inoculation increased leaf mortality compared to inoculation with *R. solani* or *C. cereale* alone by 1.9 or 4.9 times, respectively. Together, these results show that disease progression can be strongly impacted by higher-order interactions among microbial symbionts.

## Introduction

When a parasite infects a host, it interacts with the host as well as the host’s resident microbes, which can range from other parasites to mutualists [1–4]. These interactions can alter the impact of parasites on host fitness [5], with implications for host population dynamics [6]. Further, they can impact parasites themselves, with potential effects on parasite transmission [7] and parasite evolution [8,9]. While these within-host interactions among coinfecting microbes have gained much attention [10–12], most studies focus on one interaction between two microbes. However, any such pairwise interaction between two microbes is only one of many interactions among the species within a community, and we hypothesize that higher-order interactions among more than two species are also important for disease progression [13,14]. As such, focusing on pairwise interactions creates a gap in our understanding. Here, we address this gap by investigating how higher-order interactions [15] among two coinfecting parasites and a hypothesized mutualist affect disease progression.

Coinfection of a host with multiple parasite species can lead to changes in parasite growth rates, disease severity, host susceptibility, and duration of infection [16–20]. Parasites may interact antagonistically, via direct mechanisms like interference competition [21,22] or indirect mechanisms like resource competition or immune-mediated apparent competition [23,24]. Parasites can also facilitate each other by producing resources that are beneficial to other parasites [25] or indirectly, through resource- or immune-mediated interactions or other host responses or manipulations [26]. These interactions within host individuals can scale up to impact population-level processes, as parasite coinfections have major consequences for parasite transmission across a host population and epidemic dynamics [27–30].

While interactions among parasites are increasingly studied in disease ecology, we have a limited understanding of interactions between parasites and microbial mutualists [4,31]. Microbial mutualists are ubiquitous in multicellular organisms and can confer a number of benefits to their hosts [32,33]. Microbial mutualists can limit parasite colonization and growth by altering nutrient availability or competing for metabolites within a shared host [34]. They can also prime a host’s immune system, which can facilitate a rapid response to invading parasites [35]. Some microbial mutualists can affect parasite success more directly as well, by producing antimicrobial compounds that can reduce parasite growth and consequent disease severity [36,37]. Because within-host parasite accumulation is often directly or indirectly linked to between-host transmission [23,38], microbial mutualists may be important drivers of epidemiological dynamics, which can have impacts on ecosystem function [39,40]. Yet, our understanding of how microbial mutualists impact disease dynamics is mostly based on investigations of pairwise interactions.

The direction and magnitude of interactions among parasites and host-associated microbiota may depend on the sequence in which they infect host individuals [41–45]. While simultaneous coinfections can occur in nature [46], more often, microbes infect a host at different time points [41]. A microbe that infects a host can impact the availability of host resources and invoke responses from the host immune system. When microbes infect a host sequentially, these impacts on a host by the early-arriving microbe can have consequences for the colonization success and within-host growth of later-arriving microbes [47,48]. Early-infecting microbes may suppress or facilitate later-infecting microbes [41,49,50], and the effects of sequential infection can be asymmetrical and depend which microbe arrived first [51].

While the importance of within-host microbial interactions for infectious diseases is well established, the study of these interactions is typically limited to considering pairwise combinations of species (but studies investigating more than pairwise interactions include [13,52]). However, the mechanisms by which two symbionts interact may be influenced by other coinfecting symbionts, similar to indirect interactions among free-living species. For example, microbial mutualists can affect available resources within a host [53], which may modify the interaction between coinfecting parasite species that rely on these host resources [15]. Alternatively, a microbe that is a mutualist of the host could also be considered a shared enemy of parasites coinfecting the host, resulting in dynamics analogous to enemy-mediated apparent competition [54]. Investigations that consider such indirect interactions promise to more mechanistically and robustly characterize the dynamics and impacts of infectious diseases.

The detection of higher-order interactions is a challenge in ecological experiments because responses to treatments may contain both direct and indirect effects [55]. Such approaches are further complicated in the context of microbial mutualists and parasites, in which the species can be difficult to detect and isolate [56]. To address these challenges, we used a series of three controlled inoculation experiments with a grass host to disentangle (1) the interactions between a focal parasite and a microbial mutualist, (2) the interactions between the focal parasite and a second coinfecting parasite, including how those interactions are modified by infection sequence, and (3) higher-order interactions among all three symbionts and the impacts on disease progression.

## Methods

### Study System

We investigated the interactions among three fungal symbionts of the host, tall fescue (*Lolium arundinaceum*): the parasite *Colletotrichum cereale*, the parasite *Rhizoctonia solani*, and the vertically transmitted systemic endophyte, *Epichloë coenophiala.* These symbionts and host are of agricultural importance, and many potential mechanisms of within-host fungal interactions have been tested experimentally, leading to specific predictions for our system. Necrotrophs, like *R. solani,* extract resources by killing host cells. In contrast, hemibiotrophs, like *C. cereale,* initially infect hosts as a biotroph, extracting resources from living cells, a phase in which they are typically asymptomatic, then switch to a necrotrophic phase, during which infection results in disease symptoms (necrosis) on its host. In a congener of *C. cereale*, the biotrophic phase typically lasts about 48 hours [57]. Biotrophs often facilitate necrotrophs via immune-mediated crosstalk, and necrotrophs inhibit biotrophs and other necrotrophs via competition for host resources [58,59]. We expected that during its biotrophic phase, *C. cereale* would facilitate *R. solani* via immune-mediated crosstalk and during its necrotrophic phase, *C. cereale* would inhibit *R. solani* via competition for host resources. Based on this, we expected *C. cereale* to facilitate *R. solani* most when the two pathogens were inoculated simultaneously (when the early growth of *R. solani* could be facilitated by the biotrophic phase of *C. cereale*).

*E. coenophiala* is a vertically transmitted systemic fungal endophyte that is considered a mutualist under most ecological conditions [60]. While *E. coenophiala* consistently acts as a defensive mutualist with regards to herbivory, *E. coenophiala* can either facilitate or suppress infection by fungal parasites via competition for host resources and changes in host immunity, depending on parasite feeding strategy [61–64]. Additionally, *E. coenophiala* produces alkaloids and other toxins [37], and these have been shown to limit growth of fungal parasites *in vitro* [65]. Halliday et al. 2017 showed that inoculation with *E. coenophiala* inhibited *C. cereale* but had no effect on *R. solani* within a host. Given the varying effects of *E. coenophiala* on *R. solani* and *C. cereale*, we hypothesized that *E. coenophiala* would alter within-host interactions between these coinfecting parasites. Specifically, because *E. coenophiala* inhibits *C. cereale,* we hypothesized that the magnitude of any facilitation or inhibition of *R. solani* by *C. cereale* would be lower when *E. coenophiala* was present.

### Experimental Approach

We investigated the effects of within-host microbial interactions on the severity of disease caused by two fungal parasites by conducting a series of inoculation experiments in laboratory growth chambers. For detailed information on experimental approach and each experiment, refer to the supplemental information.

Each experiment involved inoculations of tall fescue plants with a different combination of three species: the endophyte, *E. coenophiala*, the fungal parasite *R. solani* and the fungal parasite *C. cereale.* To inoculate with *E. coenophiala*, plants were grown from endophyte-inoculated and endophyte-free seed. We used seed of tall fescue cultivar KY-31 provided by Dr. Tim Phillips at the University of Kentucky. The isolates of *R. solani* used in these experiments were isolated from tall fescue and phylogenetically placed within *R. solani* anastomosis group AG1-IB (sequences will be deposited in GenBank). The *C. cereale* isolate used in these experiments was also isolated from tall fescue. In addition to plants inoculated with one or both parasites, each experiment included plants that were mock-inoculated to control for cross contamination of parasites.

Following all experiments, plants were harvested. We tested for infection by the endophyte *E. coenophiala* using an immunoblot assay (Agrinostics Ltd. Co, Watkinsville, GA, USA). In all experiments, all plants grown from endophyte-free seed were confirmed after harvest to be endophyte-free; success of endophyte inoculations are reported below. Aboveground biomass was harvested, dried, and weighed.

### Inoculation experiments

#### Experiment 1

To test if and how endophyte, *E. coenophiala,* affects the within-host growth of parasite, *R. solani*, we factorially manipulated endophyte inoculation (two levels: endophyte-inoculated and endophyte-free) and *R. solani* isolate (three different isolates, plus a mock-inoculated control). The design enabled us to investigate the potential effect of the endophyte on *R. solani,* as well as if there is any parasite intraspecific variation in this effect. Plants in which the endophyte was inoculated but not detected at the conclusion of the experiment were excluded from analyses. None of the mock-inoculated control plants exhibited symptoms of *R. solani*; these plants were also excluded from analyses. The experiment began with 95 plants, and these exclusions left 67 plants in our analyses (Table S1).

#### Experiment 2

We tested how coinfection, as well as infection sequence in coinfection, with the parasite *C. cereale*, affects the severity of disease caused by *R. solani.* Specifically, we inoculated endophyte-free plants with *C. cereale* alone, *R. solani* alone, or with both parasites. We had three treatments in which both parasites were inoculated: Simultaneous co-inoculation, sequential inoculation in which *C. cereale* was inoculated first, and sequential inoculation in which *R. solani* was inoculated first. None of the mock-inoculated control plants exhibited symptoms of either parasite; these plants were excluded from analyses. The experiment began with 60 plants, and these exclusions left 50 plants in our analyses (Table S2).

#### Experiment 3

To test if endophyte, *E. coenophiala,* shifts competitive outcomes of two coinfecting parasites, we factorially manipulated inoculation with *E. coenophiala*, *C. cereale,* and *R. solani*. In treatments in which both parasites were inoculated, parasites were inoculated simultaneously because as further discussed below, in Experiment 2, *C. cereale* had the most impact on *R. solani* within-host growth when the parasites were inoculated simultaneously. Plants in which the endophyte was inoculated but not detected at the conclusion of the experiment were excluded from analyses. None of the mock-inoculated control plants exhibited symptoms of either parasite; these plants were also excluded from analyses. The experiment began with 122 plants, and these exclusions left 91 plants in our analyses (Table S3).

### Data Analysis

Each model analyzed one experiment and included one dependent variable pertaining to one parasite species. Individual leaves were analyzed as hosts because each parasite infection is restricted to a single leaf. We analyzed all data in R v. 3.6.1 (R Core Team, 2017).

To evaluate within-host interactions that determine *R. solani* growth rate, we modelled *R. solani* disease severity as a linear function of days after inoculation, parasite treatment (or isolate, in the case of Experiment 1), endophyte treatment (in Experiments 1 and 3) and all interactions as fixed effects. Additionally, we included an interaction between chamber and shelf as a fixed effect to account for any variation due to location of plants. We included random intercepts for leaf ID and by-leaf random slopes for days after *R. solani* inoculation to account for longitudinal surveying. In Experiment 2, two leaves per plant were inoculated and surveyed, so in models analyzing this data, we nested the random intercepts for leaf ID and the by-leaf random slopes for days after *R. solani* inoculation within plant ID. All models of parasite growth were implemented using the lme function in the nlme package (version 3.1.142, [67]).

To evaluate within-host interactions that determine *C. cereale* symptom development (in Experiment 3), we performed a survival analysis using Cox proportional hazards models to quantify the instantaneous risk over time of a leaf developing symptoms of *C. cereale* (using the coxph function within the survival package (version 2.43, [68]). In each survival model, we accounted for placement within the growth chambers (chamber and shelf) using the cluster function within the survival package.

To test how parasite inoculation treatment, endophyte treatment and their interaction affect host mortality (in Experiment 3), we performed three survival analyses with leaf survival as the response variable. First, to consider how each parasite species and their interaction affected leaf survival, we conducted a survival analysis with parasite inoculation treatment as a predictor variable (4 levels: mock-inoculated control, *C. cereale* alone, *R. solani* alone, and coinfection). Second, to consider whether the impact of *C. cereale* on survival of leaves infected by *R. solani* could be modified by the endophyte, *E. coenophiala*, we conducted a survival analysis including the endophyte treatment, a subset of the parasite treatment levels (single- or co-inoculation, *i.e.* only *R. solani* or both *R. solani* and *C. cereale*), and their interaction. Third, to consider whether the impact of *R. solani* on survival of leaves infected by *C. cereale* could be modified by the endophyte, *E. coenophiala*, we conducted a survival analysis including the endophyte treatment, a subset of the parasite treatment levels (single- or co-inoculation, *i.e.* only *C. cereale* or both *R. solani* and *C. cereale*), and their interaction. All analyses used Cox proportional hazards models to quantify the instantaneous risk over time that a leaf would die. In each model, we accounted for placement within the growth chambers (chamber and shelf) using the cluster function. Pairwise comparisons among treatments were made by releveling the parasite treatment predictor to change the reference level. Disease severity can reduce host survival, so to evaluate if effects of inoculation treatments on leaf survival were explained by disease severity, we expanded upon each model by adding a time-dependent covariate of disease severity as estimated by percent leaf damaged.

To evaluate how within-host microbial interactions impact host growth (in Experiment 3), we modelled host aboveground biomass at the end of the experiment as a linear function of endophyte treatment (2 levels: Endophyte Absent and Endophyte Infected), parasite inoculation treatment (3 levels: *C. cereale* alone, *R. solani* alone, and Coinfection), and their interaction. Additionally, we included an interaction between chamber and shelf as a fixed effect to account for any variation due to location of plants. We used emmeans ([69], version 1.3.2) to evaluate the estimated marginal means of host aboveground biomass for each explanatory variable level, adjusted for multiple comparisons (Tukey HSD).

To improve statistical inference, instead of emphasizing statistically significant results, we report our results using the language of “statistical clarity” [70].

## Results

### How is disease severity of *R. solani* affected by a systemic endophyte?

Experiment 1 factorially manipulated inoculation with the endophyte *E. coenophiala,* and inoculation with three isolates of the parasite *R. solani*. There was no clear effect of endophyte infection on the severity of disease caused by *R. solani*, as estimated by *R. solani* lesion length, or by change in lesion length over time. While *R. solani* disease severity increased over time (p<0.0001), endophyte infection had no clear main effect (p = 0.13) and no clear interactive effect with days after *R. solani* inoculation (p = 0.59) on *R. solani* lesion length over time (Figure 1, Table S4). While isolate of *R. solani* had a clear main effect on disease severity (p < 0.0001), there was no evidence of disease caused by different isolates progressing at different rates (p = 0.60). Overall, accounting for variation among isolates of *R. solani,* endophyte infection had no clear effect on progression of disease caused by *R. solani* within a shared host.

**Figure 1:**
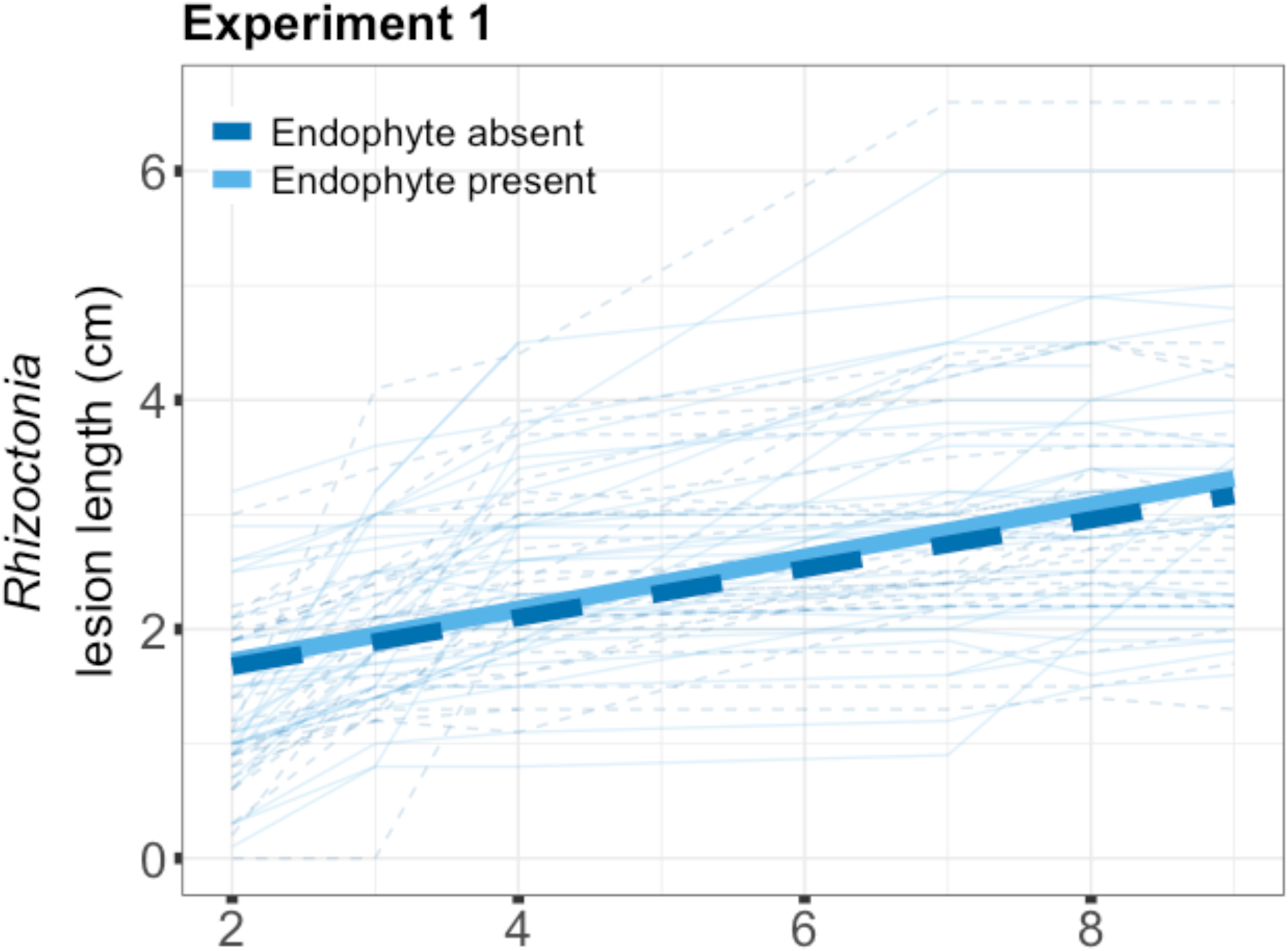
In Experiment 1, *E. coenophiala* had no clear effect on severity of disease caused by *R. solani*. Disease severity was quantified by *R. solani* lesion length. The faint lines are individual leaves and bolded lines are the output of a linear mixed effects model incorporating endophyte infection and days after inoculation with *R. solani*. Lesion length clearly increased over time, but there was no clear effect of the presence of the endophyte on *R. solani* growth rate.

### How is disease severity of *R. solani* affected by coinfection with the parasite *C. cereale*?

Experiment 2 manipulated inoculation sequence of the parasite *C. cereale* relative to *R. solani*, in the absence of the endophyte *E. coenophiala.* The severity of disease caused by *R. solani* (lesion length) increased over time (p<0.0001), and that rate of increase depended on the inoculation sequence of *R. solani* relative to *C. cereale* (Treatment : Days After Inoculation, p=0.022, Figure 2, Table S5.A). Specifically, the increase over time of *R. solani* disease severity was more than twice as high in the treatments in which *C. cereale* was inoculated before or at the same time as *R. solani* than the treatment in which *R. solani* was inoculated on its own (p = 0.05, Table S5.B), indicating a facilitative effect of *C. cereale* on *R. solani*. When *R. solani* was inoculated before *C. cereale,* the increase over time in *R. solani* disease severity was 1.38 times higher than it was when *R. solani* was inoculated on its own, but this increase lacked statistical support (p = 0.83, Table S5.B). Together, these results suggest that the increase over time in *R. solani* disease severity is higher when in coinfection with *C. cereale,* but the magnitude of this effect depends on the infection sequence of the parasites.

**Figure 2:**
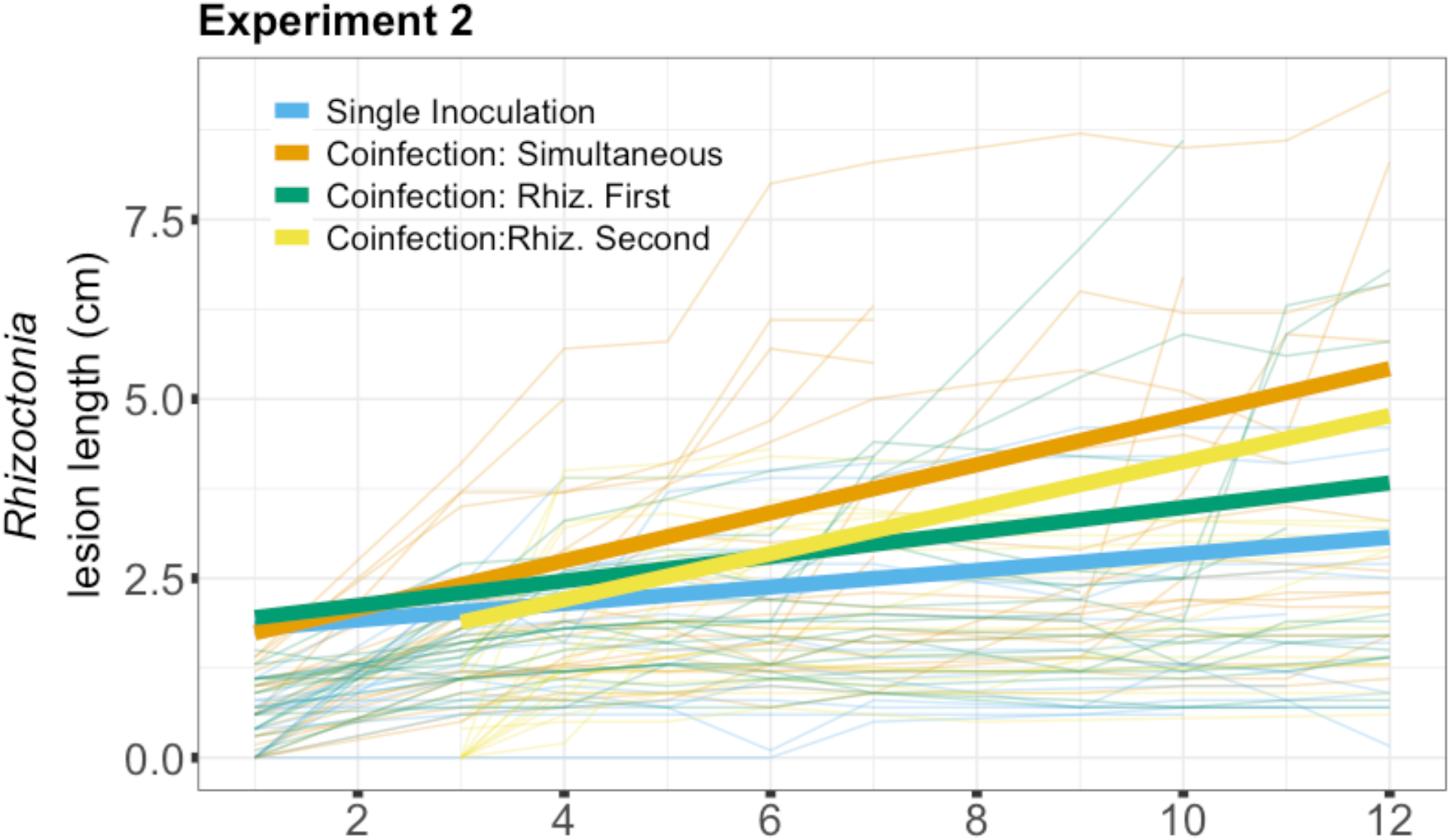
In Experiment 2, the severity of disease caused by *R. solani* was higher when in coinfection with *C. cereale* than when inoculated on its own. Disease severity was quantified as *R. solani* lesion length. Each faint line represents a leaf, and bolded lines are model output from a linear mixed effects model than incorporated parasite treatment and time after inoculation with *R. solani*. *R. solani* growth rate was higher in the coinfection treatments in which *C. cereale* and *R. solani* were inoculated simultaneously and when *R. solani* was inoculated after *C. cereale* than when *R. solani* was inoculated on its own (p ≤ 0.05).

### How is disease severity of *R. solani* affected by the interaction between a fungal parasite, *C. cereale,* and a systemic endophyte?

Experiment 3 factorially manipulated inoculation of the endophyte *E. coenophiala* and the parasites *C. cereale* and *R. solani*; the parasite inoculations were simultaneous. Consistent with the results of Experiment 2, progression of disease caused by *R. solani* was facilitated by co-inoculation with *C. cereale*. Specifically, the slope of *R. solani* lesion length over time was 2.26 times higher for leaves co-inoculated with *C. cereale* than leaves inoculated with only *R. solani* (Figure 3, Table S6, Parasite Treatment : Days After Inoculation, p = 0.005). Severity of disease caused by *R. solani* tended to progress more quickly when co-inoculated with *C. cereale* in plants infected with the endophyte, relative to plants that were endophyte-free, but this three-way interaction had weak statistical support (Table S6, p = 0.11).

**Figure 3:**
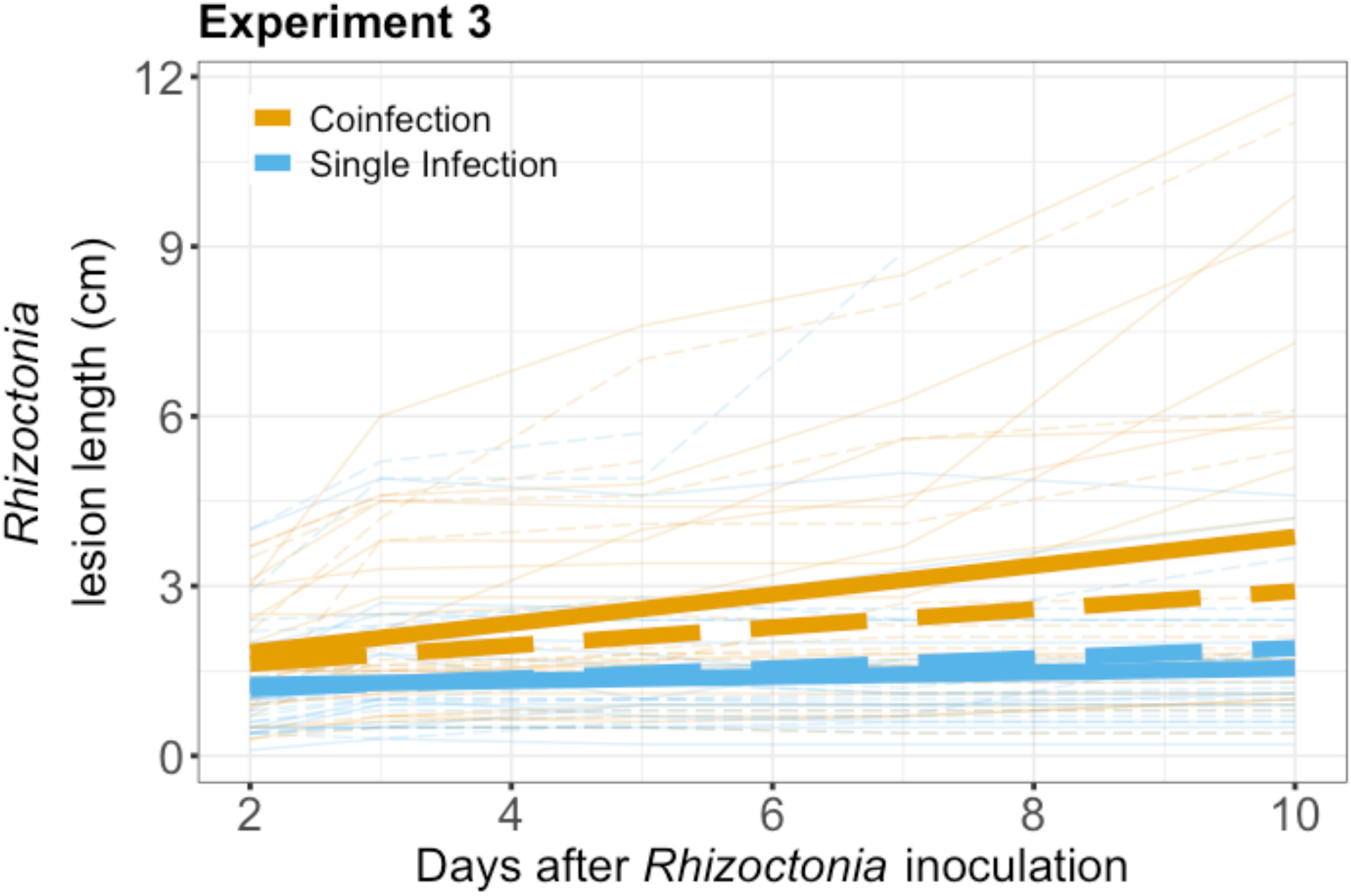
In Experiment 3, the growth rate of *R. solani* was higher when in coinfection with *C. cereale* than when inoculated on its own. *R. solani* lesion length over time. Faint lines are individual leaves, and bold lines are model output from a linear mixed effects model that incorporated parasite treatment and time after inoculation with *R. solani*. There was an interaction between time after inoculation and parasite treatment (2 levels: single infection vs. coinfection). Additionally, *R. solani* growth rate tended to increase even more under coinfection with *C. cereale* when the endophyte was also present (solid lines) than when it was absent (dashed lines), but this had weak statistical support.

We also quantified how *C. cereale* responded to *R. solani* and *E. coenophiala. C. cereale* is a hemibiotroph, initially infecting leaves asymptomatically as a biotroph and then shifting to a symptomatic necrotrophic phase. We considered if and how coinfection with *R. solani* and *E. coenophiala* affected the development of *C. cereale* symptoms (here, the development of chlorosis). We conducted a survival analysis investigating factors that affected the time until *C. cereale* symptoms were exhibited. Compared to leaves that were only inoculated with *C. cereale*, leaves inoculated simultaneously with both *C. cereale* and *R. solani* had 0.42 times the hazard of developing *C. cereale* symptoms (Figure S1, Table S7.A, p<0.001). Thus, *R. solani* infection slowed the development of *C. cereale* symptoms. By the end of the experiment, 73% of leaves inoculated with *C. cereale* alone exhibited symptoms of the parasite, compared to 33% of leaves that were co-inoculated and infected with *R. solani*. While this impact of *R. solani* on *C. cereale* symptom development held in the presence and absence of *E. coenophiala,* there was less statistical support for *R. solani* impacting *C. cereale* symptom development in endophyte-infected plants (Table S7.B, p = 0.0870). Over the course of the experiment, *E. coenophiala* had no clear direct impact on *C. cereale* symptom development (Table S7.A, p = 0.516).

In addition to impacting parasite lesion growth and disease symptoms, within-host microbial interactions also impacted host individuals in terms of leaf mortality and plant biomass. Leaves that were inoculated with both parasites had greater mortality than leaves in all other parasite inoculation treatments (Figure 4A, Table S8.A, Wald statistic=12.34 on 3df, p = 0.006). Compared to mock-inoculated control leaves, leaves inoculated with only *C. cereale* had a 3.2 times higher hazard (p = 0.08), leaves inoculated with only *R. solani* had a 5.2 times higher hazard (p = 0.03), and leaves inoculated with both parasites had an 8.0 times higher hazard (p = 0.01) of mortality.

**Figure 4:**
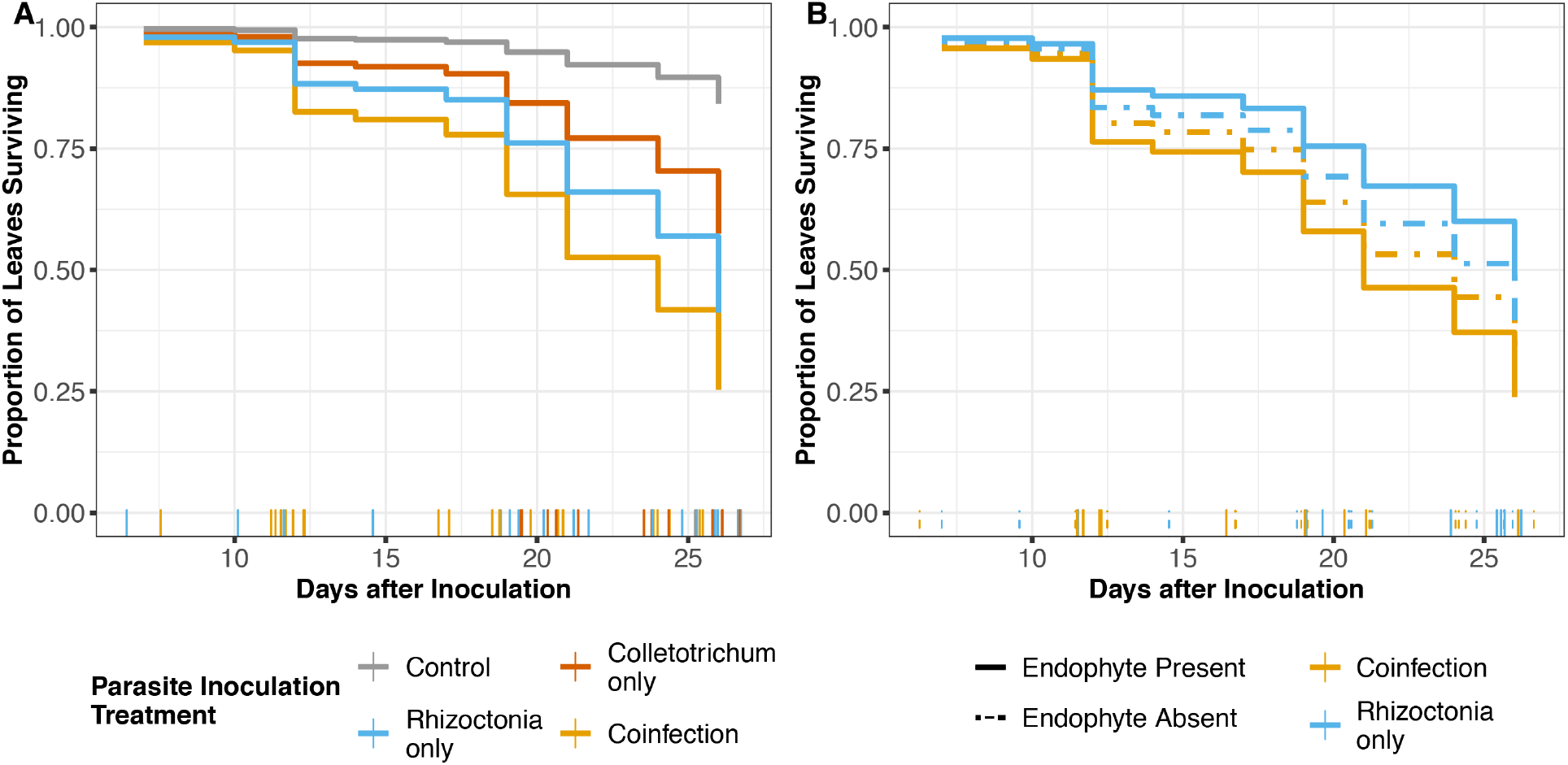
Parasite co-inoculation increased leaf mortality. Model-estimated relative risk of leaf death over time. Plots are results of Cox proportional hazards models. For leaves that died, vertical lines along the x-axis show when (in days after inoculation) each leaf died. X-axis values of those vertical lines are jittered to show the data. A. Parasite treatment affected leaf mortality. Compared to leaves that were mock-inoculated, leaves inoculated with only *C. cereale* had 3.22 times higher hazard of mortality, though with weak statistical support (p = 0.08), leaves inoculated with only *R. solani* had 5.17 times higher hazard of mortality (p = 0.03), and leaves that were co-inoculated with both parasites had an 8.00 times higher hazard of mortality (p = 0.01). B. While there was no difference in leaf mortality between leaves inoculated with just *R. solani* and leaves co-inoculated with *R. solani* and *C. cereale* in endophyte-free plants (p = 0.6), leaves co-inoculated with both parasites had a 1.94 times higher hazard of mortality than leaves inoculated with only *R. solani* in endophyte-infected plants (p = 0.01).

To test whether *C. cereale* inoculation and endophyte infection interacted to influence mortality of leaves infected with *R. solani*, we used a second Cox proportional hazards model that included a four-level predictor variable representing the four factorial combinations of parasite treatment levels that involved *R. solani* (2 levels: inoculation with only *R. solani* and co-inoculation with *C. cereale*) and endophyte infection (2 levels: free or infected). The effect of parasite co-inoculation on leaf mortality depended on endophyte infection (Wald statistic=8.59 on 3df, p = 0.04). While there was no clear difference in leaf mortality between leaves inoculated with only *R. solani* and leaves co-inoculated with *R. solani* and *C. cereale* in endophyte-free plants (p = 0.581), parasite co-inoculation increased leaf mortality in endophyte-infected plants, with a 1.9 times higher hazard than for leaves inoculated with only *R. solani* (Figure 4B, Table S8.B, p = 0.012). To test if the interaction between *R. solani* and endophyte infection could similarly influence mortality of leaves infected with *C. cereale,* we used a third Cox proportional hazards model that was similarly structured, with a four-level predictor variable representing the four factorial combinations of parasite treatment levels that involved *C. cereale* (2 levels: inoculation with only *C. cereale* and co-inoculation with *C. cereale*). Similar to the *R. solani* results, while there was no clear difference in leaf mortality between leaves inoculated with only *C. cereale* and leaves co-inoculated with *R. solani* and *C. cereale* in endophyte-free plants (p = 0.180), parasite co-inoculation increased leaf mortality in endophyte-infected plants, with a 4.9 times higher hazard than for leaves inoculated with only *C. cereale* (Figure S2, Table S8.C, p = 0.008).

Leaf mortality increased not only with parasite inoculation treatment, but with disease severity, as estimated by overall percent leaf area damaged (Table S9.A,B,C p<0.001). Including total disease severity as a time-dependent covariate in the survival analyses largely accounted for the impacts of parasite inoculation treatments (Table S9.A). Disease severity accounted for the interactive effect of *C. cereale* and endophyte treatment on mortality of leaves infected with *R. solani* (Table S9.B), but did not account for the interactive effect of *R. solani* and endophyte treatment on mortality of leaves infected with *C. cereale* (Table S9.C). These results suggest that differences in leaf mortality among parasite inoculation treatments were largely through effects on disease severity.

At the end of the experiment, plants tended to have the lowest biomass when coinfected with *E. coenophiala, C. cereale,* and *R. solani*. Across parasite treatments, plants infected with the endophyte *E. coenophiala* produced less biomass than plants lacking the endophyte (Table S10, S11, p=0.02). While the interaction between *E. coenophiala* and parasite treatment was not supported by the ANOVA test (Table S10, p=0.20), Tukey HSD indicated that plants inoculated with all three fungal species may have produced the lowest biomass. Specifically (Figure S3, Table S11), plants inoculated with all three fungal species tended to have less final biomass than plants inoculated with just *C. cereale* (Tukey HSD, p = 0.055), plants inoculated with just *R. solani* (Tukey HSD, p = 0.052), and plants inoculated with both *C. cereale* and *R. solani* (Tukey HSD, p = 0.075). While these differences were not statistically clear, together they suggest that plants coinfected with the endophyte and both parasites produced the least biomass.

## Discussion

In this study, interactions among a microbial mutualist and two coinfecting parasites altered disease progression. The growth rate of *R. solani* within a host was higher when in coinfection with the parasite *C. cereale*, which was observed in both experiments that tested this interaction (Experiments 2 and 3). *R. solani* slowed the development of *C. cereale* symptoms (Experiment 3). While there was no direct effect of systemic endophyte, *E. coenophiala,* on *R. solani* lesion growth (Experiment 1) or on *C. cereale* symptom development (Experiment 3), the magnitude of the effect of *C. cereale* on *R. solani* tended to be greater when *E. coenophiala* was also present (Experiment 3), though this trend had weak statistical support. Further, leaf mortality was only affected by parasite coinfection when *E. coenophiala* was present. Specifically, when *E. coenophiala* was present, leaf mortality was greater when *C. cereale* and *R. solani* were in coinfection compared to when either was inoculated on its own. Further, plants tended to have the lowest biomass when coinfected with *E. coenophiala, C. cereale,* and *R. solani*. Together, these results show that higher-order interactions between microbial symbionts can have important impacts on both symbionts and hosts.

The presence of the systemic endophyte, *E. coenophiala,* had no effect on within-host growth of the parasite *R. solani* in the absence of *C. cereale*. While systemic grass endophyte infection has been shown to protect host plants by increasing resistance to herbivores and seed predators, as well as protect against abiotic stressors [71,72], evidence for defending against infectious disease is less consistent. There are many studies in which such systemic grass endophytes limit susceptibility to and progression of diseases caused by certain parasites [73–75], but there are also studies that document no effect of these endophytes on parasites and the development of disease symptoms [76,77], and there are still other studies that report higher susceptibility to parasites when grass hosts are infected with an endophyte [78]. In another study that included *E. coenophiala* endophyte infection of tall fescue, the endophyte limited progression of disease symptoms caused by *R. solani*’s congener *R. zeae* in most host genotypes, but had no effect in some genotypes [65]. This literature and our results suggest that the relationship between systemic grass endophytes and infectious disease is not easily generalizable, and the interactions between these endophytes and parasites may depend on parasite species, host genotype, and environmental conditions.

While *E. coenophiala* had no direct effect on *R. solani, C. cereale* increased the within-host growth of *R. solani*. Further, the magnitude of this facilitation depended on infection sequence. There is growing evidence that infection sequence can determine the magnitude and direction of the effect of within-host microbial interactions [42,44,45]. The effect of infection sequence on microbial interactions may depend on the microbes’ feeding modes, particularly whether or when they feed on living or dead plant tissue.

*Rhizoctonia solani* is a necrotroph, and extracts resources by killing host cells. *C. cereale* is a hemibiotroph, with an initial asymptomatic phase in which it extracts resources as a biotroph, followed by a symptomatic phase in which it extracts resources as a necrotroph. Host plants respond to biotrophs and necrotrophs via the salicylic acid and jasmonic acid resistance pathways, respectively, so infection by a microbe with either feeding mode can trigger cross-resistance against other microbes with that same feeding mode. Furthermore, crosstalk between these two pathways can result in immune-mediated interactions between coinfecting parasites of contrasting feeding modes; for example, a microbe that induces the salicylic acid pathway can thereby indirectly suppress the jasmonic acid pathway [24]. Via such crosstalk, biotrophs are expected to facilitate necrotrophs, while via cross-resistance, necrotrophs are expected to inhibit other necrotrophs [58,59]. In our Experiment 2, the greatest increase in the necrotroph *R. solani*’s growth rate occurred when *C. cereale* and *R. solani* were inoculated simultaneously, which would be when C. cereale was in biotrophic mode. Our results are therefore consistent with the initial biotrophic phase of *C. cereale* facilitating the growth of *R. solani* via crosstalk between the host immune pathways.

The facilitative effect that *C. cereale* had on *R. solani* was greatest on plants that were also infected with the endophyte *E. coenophiala.* Additionally, endophyte-infected plants’ leaves that were co-inoculated with both fungal parasites had greater mortality and endophyte-infected plants that were co-inoculated with both fungal parasites had lower aboveground biomass. While *E. coenophiala* did not have a direct effect on how disease caused by *R. solani* progressed within a host or impacted leaf mortality, our experiments indicated that *E. coenophiala* impacted *R. solani* indirectly. The two canonical types of indirect effects are interaction chains and interaction modifications [15]. An interaction chain is when two species are linked to changes in the abundance of a third species, whereas an interaction modification results when a third species alters the nature of an interaction between two other species. For example, if one species alters the phenotype of a second species, the trait change in the second species can alter its effect on a third species [79,80]. In our experiments, *E. coenophiala* had no clear impact on *C. cereale* symptom development, so if symptom development correlates with growth of *C. cereale* within the host, then our results would be consistent with an interaction modification. Such an interaction modification might be mediated by traits such as the timing of *C. cereale*’s switch from a biotrophic to a necrotrophic feeding strategy, but testing that mechanism would require further experimentation. Both Abbate et al. (2018) and Marchetto & Power (2017) suggest that the outcome of interactions between two parasite species can be impacted by a third symbiont, whether that third symbiont be a third parasite in the case of Abbate et al. (2018) or a microbial mutualist in the case of Marchetto & Power (2017). Our results provide further evidence that higher-order interactions between microbial symbionts involving two or more parasites can impact disease progression, advancing on these recent studies by demonstrating that these interactions can have particularly strong consequences for host mortality.

This study makes clear that higher-order interactions can alter disease progression. As these complex interactions are challenging to probe under field conditions, we posit that indirect interactions may drive differences in the magnitude and direction of within-host microbial interactions in lab- vs. field-based studies. There is mounting evidence that effects of microbial interactions under lab settings may not scale up to those under field conditions [81]. In contrast to the facilitation of *R. solani* by *C. cereale* presented here based on lab inoculations, Halliday et al. (2017) documented infections from ambient inoculum in the field that indicated an antagonistic interaction between *C. cereale* and *R. solani.* It may be that a limitation to directly extrapolating findings from lab experiments to field settings is the presence of higher-order interactions among many symbionts that are difficult to detect. Future work that considers higher-order interactions under field settings may clarify this connection.

We found that a microbial mutualist affected the interaction between two coinfecting parasites, having effects on both the parasites and the host. As coinfections have gained interest within disease ecology, studies of pairwise interactions between coinfecting microbes have increased, but higher-order interactions among more than two coinfecting microbes are still underexplored. Our results suggest that pairwise interactions may be context-dependent. The outcomes of pairwise interactions may depend on the presence of other species in the community. These higher-order interactions have important impacts on both hosts and symbionts and may be a mechanism that explains why results of pairwise interaction studies in the lab are not always consistent with those in the field. Future research should therefore consider the larger microbial community associated with a host when evaluating the impacts of coinfections.

## Acknowledgements

We thank Marc Cubeta for helpful conversations about the biology of *Rhizoctonia* and James Umbanhowar for helpful comments and help with statistics and Brooklynn Joyner for help with data collection. We thank Tim Phillips for the tall fescue seed used for experiments and Fletcher Halliday for providing the isolate of *C. cereale* used in inoculations. This work was supported by the NSF-USDA joint program in Ecology and Evolution of Infectious Diseases (USDA-NIFA AFRI grant 2016-67013-25762). K.R.O. was supported by graduate research fellowships from the Triangle Center for Evolutionary Medicine and the National Science Foundation.

## Notes

### Competing Interest Statement

The authors have declared no competing interest.

