## Supplementary Material for "Higher-order interactions among coinfecting parasites and a microbial mutualist impact disease progression"

Supplemental Methods

Experimental Approach

We investigated the effects of within-host microbial interactions on the disease outcomes caused by two fungal parasites by conducting a series of three inoculation experiments in laboratory growth chambers. For Experiment 1 and 2, plants were grown in 993 mL pots (D60 Deepots, Stuewe and Sons, Corvallis OR), and for Experiment 3, plants were grown in 656 mL pots (D40 Deepots, Stuewe and Sons, Corvallis OR). In all experiments, plants were grown in MetroMix 360 potting mix (Sun Gro Horticulture, Agawam, MA, USA). The plants were watered from the bottom of the pot during the experiments. The experiments were conducted in two growth chambers with a 12-hour dark/12-hour light schedule and kept at 29°C. Humidifiers were placed on each shelf of each growth chamber and used to maintain relative humidity at 85-90%. The locations of individual plants of each treatment were randomized within each shelf of each growth chamber.

Prior to each experiment, we started plants from seed in a greenhouse. The greenhouse temperature was kept between 19.7-22.2ºC and light was supplemented between 9am and 7pm if natural light fell below 350 W/m$^2$. Plants were relocated from the greenhouse to growth chambers (Percival PGC-6L; Perry, Iowa) six weeks after germination and given two days to acclimate before starting an experiment.

Each experiment involved a different combination of inoculations with endophyte, *R. solani* and/or *C. cereale.* We used endophyte-inoculated and endophyte-free seed provided by Dr. Tim Phillips at the University of Kentucky.

Inoculations with *R. solani* involved placing a 0.6 cm PDA plug covered with actively growing mycelia at the base of the oldest leaf of a random tiller of a plant. Successful infection of *R. solani* requires that the inoculation site be humid. To keep the inoculation site humid, we covered the PDA plug on the leaf with a piece of cotton wet with sterile water, and wrapped the base of the leaf with tin foil, which was then wrapped in parafilm. The plug, cotton, foil and parafilm were removed after two days. The *R. solani* strains used in these experiments were isolated from tall fescue in Duke Forest Research and Teaching Laboratory in summer 2015.

Inoculations with *C. cereale* occurred in one of two ways. In Experiment 2, we inoculated with mycelia in the same way as described above for *R. solani.* In Experiment 3, we inoculated with *C. cereale* spores (as described in Beirn, Wang, Clarke, & Crouch, 2015). *C. cereale* was grown in culture on PDA plates under constant light for 10 days until the *C. cereale* had filled the plates. Under sterile conditions, plates were each flooded with 10 milliliters of sterile water and a sterile loop was used to scrape the spores from the original culture. Then, the sterile loop was used to smear a new plate with spore solution. These plates were grown under constant light without being sealed for 4 days. After four days, these plates were flooded again and the resulting solution was saved. Spore concentration was adjusted to $10^6$ spores/mL and 10% potato dextrose broth (PDB). To inoculate plants, 10 milliliters of spore solution were sprayed onto the plant using an atomizer, and to keep the conditions humid, the plants were covered with a plastic bag for two days. The *C. cereale* strain used in these experiments was isolated from tall fescue in the Duke Forest Research and Teaching Laboratory in the fall of 2015 [2].

Within each experiment, we also included mock-inoculated controls. When the inoculation involved PDA plugs, mock-inoculated plants received sterile PDA plugs. When the inoculation involved a spore suspension in 10% PDB, mock-inoculated plants were sprayed with sterile 10% PDB. In all other ways, mock-inoculated plants were treated the same as inoculated plants.

Following all experiments, plants were harvested. For experiments in which the endophyte was involved, a one-inch-long cross-section from the base of each of two tillers per plant was collected and frozen. The cross-sections were used to confirm the presence of the endophyte in aboveground tissue with an immunoblot assay (Agrinostics Ltd. Co, Watkinsville, GA, USA).

Inoculation experiments

*Experiment 1*

To test if and how the systemic endophyte, *E. coenophiala,* affects the within-host growth and disease severity of *R. solani*, we conducted an inoculation experiment in which we factorially manipulated endophyte inoculation (two levels: inoculated and mock-inoculated) and *R. solani* strain (three strains and mock inoculation). The design enabled us to investigate the potential effect of the endophyte on *R. solani,* as well as parasite intraspecific variation in this effect. Plants within each endophyte treatment (endophyte-inoculated and endophyte-free) were fully randomized within 4 parasite treatments: inoculation with one of three strains of *R. solani* or mock inoculation. Following inoculations, the presence/absence of disease symptoms, length of lesions, and survival of leaves were observed and measured. Individual leaves were tracked longitudinally, with multiple observations per leaf over time. Observations were made on

6 days (2, 3, 4, 7, 8, and 9 days after inoculation with *R. solani*). In total, across the 6 observation days and 67 plants included in analyses (Table 1), there were 402 potential observations, but leaf mortality precluded 30 potential observations, leaving 372 total observations in our analysis.

To evaluate how endophyte infection determines *R. solani* growth rate, we modelled *R. solani* disease severity (mean-centered to aid in statistical interpretation) as a linear function of endophyte infection, *R. solani* strain, time after *R. solani* inoculation, and all interactions as fixed effects. The model had the following general form:

**Fixed effects:** Chamber:Shelf **+** cDAI * E * Strain
**Random effects**: Random intercepts for Leaf ID, and by-host random slopes for cDAI

where cDAI represents the centered time after *R. solani* inoculation, E represents the infection status by the endophyte (0,1), Strain represents the *R. solani* strain (1, 2, or 3), Chamber represents growth chamber location (0,1) and Shelf represents the shelf within the growth chamber (Top, Bottom).

*Experiment 2*

To test how coinfection, as well as infection sequence, with a parasite, *C. cereale*, affects the severity of disease caused by *R. solani,* we conducted an inoculation experiment with the two parasites (all plants were endophyte-free). Specifically, we inoculated two leaves per plant with *C. cereale* alone, *R. solani* alone, or both parasites. We had three treatments in which both parasites were inoculated: Simultaneous co-inoculation, sequential inoculation in which *C. cereale* was inoculated first, and sequential inoculation in which *R. solani* was inoculated first. For the sequential inoculations, the second inoculation occurred when symptoms of the first parasite appeared on all leaves in the relevant treatment. It took 10 days for *C. cereale* symptoms (chlorosis) to be observed on all inoculated leaves and 2 days for *R. solani* symptoms (necrosis) to be observed on all inoculated leaves. Inoculation of *C. cereale* was performed with mycelia on PDA plugs as described above.

Following inoculations, symptoms of *R. solani* were measured as lesion length. We did not analyze symptoms of *C. cereale* as a response because it did not produce necrotic lesions in this experiment, perhaps owing to being inoculated with mycelium instead of spores as in Experiment 3. Individual leaves were tracked longitudinally, with multiple observations per leaf over time. Observations were made on 10 days (1, 3, 4, 5, 6, 7, 9, 10, 11, and 12 days after inoculation with *R. solani*), except that leaves inoculated first with *C. cereale* then second with *R. solani* were observed on 9 days (3, 4, 5, 6, 7, 8, 10, 11, and 12 days after inoculation with *R. solani*). In total, across the 9 or 10 observation days, 2 inoculated leaves per plant, and 50 plants included in analyses (Table 2), there were 980 potential observations, but leaf mortality precluded 295 potential observations, leaving 685 total observations in our analysis.

To evaluate how coinfection and infection sequence determine *R. solani* growth rate, we modelled *R. solani* disease severity (mean-centered to aid in statistical interpretation) as a linear function of parasite inoculation treatment, time after inoculation, and all interactions. The model had the following general form:

**Fixed effects:** Chamber:Shelf **+** cDAI * Parasite_Treatment
**Random effects**: Random intercepts for Leaf ID nested within Plant ID, and by-host random slopes for cDAI

where cDAI represents the centered time after *R. solani* inoculation, Parasite_Treatment represents the experimental inoculation treatment (4 levels: single inoculation of focal parasite, coinfection: simultaneous inoculation, coinfection: focal parasite inoculated first, and coinfection:focal parasite inoculated second), Chamber represents growth chamber location (0,1) and Shelf represents the shelf within the growth chamber (Top, Bottom).

*Experiment 3*

To test if and how a systemic endophyte will shift competitive outcomes of two coinfecting parasites, we conducted an inoculation experiment in which we factorially manipulated inoculation with *E. coenophiala*, *C. cereale,* and *R. solani*. In the coinfection treatment, both parasites were inoculated simultaneously because as further discussed below, in Experiment 2, *C. cereale* had the most impact on *R. solani* within-host growth when the parasites were inoculated simultaneously. Inoculum of *C. cereale* was in the form of a spore suspension, as discussed above.

Following inoculations, leaves were surveyed for disease severity. Individual leaves were tracked longitudinally, with multiple observations per leaf over time. The length of each *R. solani* lesion was measured on 5 days (2, 3, 5, 7, and 10 days after inoculation with *R. solani*). In total, across the 5 observation days and 91 plants included in analyses (Table 3), there were 455 potential observations, but leaf mortality precluded 177 potential observations, leaving 278 total observations in our analysis of *R. solani* lesion length. Three times per week throughout the experiment, we also estimated the percent of leaf area infected with each parasite by visually comparing leaves to reference images of leaves of known percent area infected (Mitchell et al. 2002, 2003; Halliday et al. 2017). Symptoms of *C. cereale* included chlorosis and small brown necrotic lesions. 26 days after inoculation, plants were harvested. Aboveground biomass was harvested, dried, and weighed.

To evaluate how endophyte infection and parasite coinfection determine *R. solani* growth rate, we modelled *R. solani* disease severity (mean-centered to aid in statistical interpretation) as a linear function of endophyte infection, parasite inoculation treatment, time after parasite inoculation, and all interactions as fixed effects. The model had the following general form:

**Fixed effects:** Chamber:Shelf **+** cDAI * Parasite_Treatment * E
**Random effects**: Random intercepts of Leaf ID, and by-host random slopes for cDAI

where cDAI represents the centered time after *R. solani* inoculation, Parasite_Treatment represents the experimental inoculation treatment (2 levels: single inoculation of focal parasite, coinfection), E represents the infection status by the endophyte (0,1), Chamber represents growth chamber location (0,1) and Shelf represents the shelf within the growth chamber (Top, Bottom).

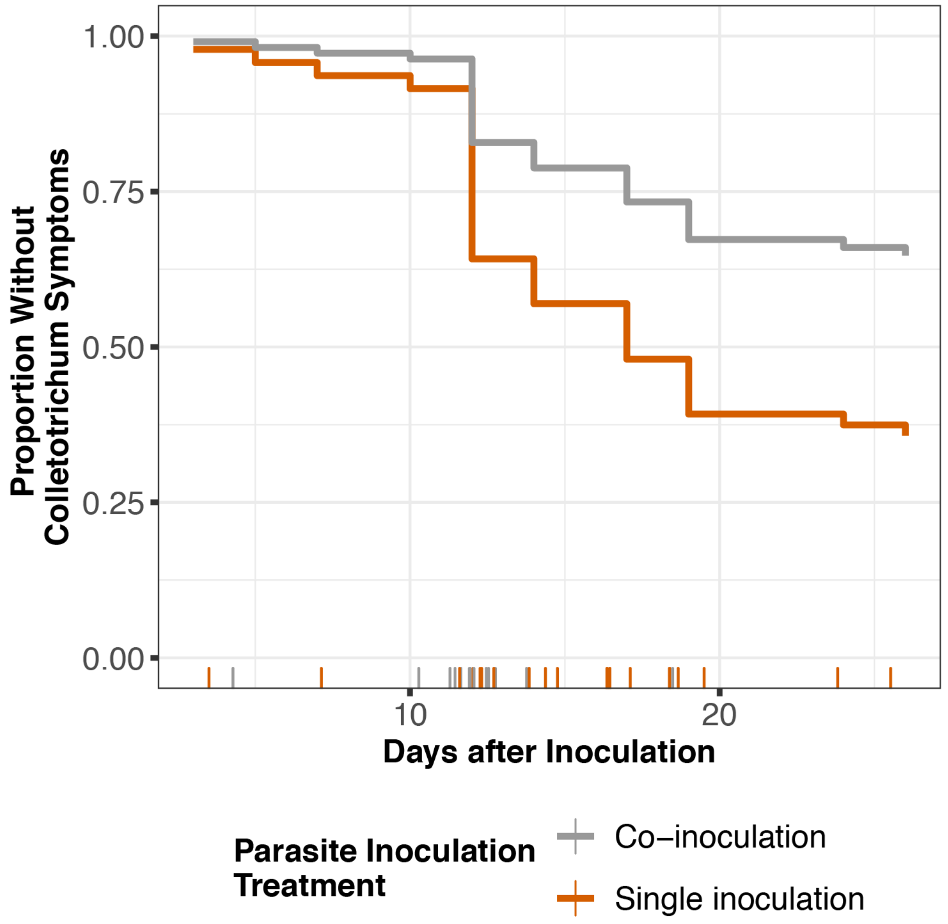


**Figure S1: Experiment 3 *C. cereale* results.** Model-estimated relative risk of *C. cereale* symptom development over time, in plants inoculated with *C. cereale* only or co-inoculated with *R. solani*. Plots are results of Cox proportional hazards models. For leaves that developed *Colletotrichum* symptoms, vertical lines along the x-axis show when (in days after inoculation) symptoms were first observed, colored by parasite inoculation treatment. X-axis values of those vertical lines are jittered to show the data. Parasite treatment clearly affected the time until *C. cereale* symptoms were exhibited. Compared to leaves that were inoculated with only *C. cereale*, leaves that were also inoculated with *R. solani* had 0.42 times the hazard of exhibiting *C. cereale* symptoms ($p < 0.0001$).

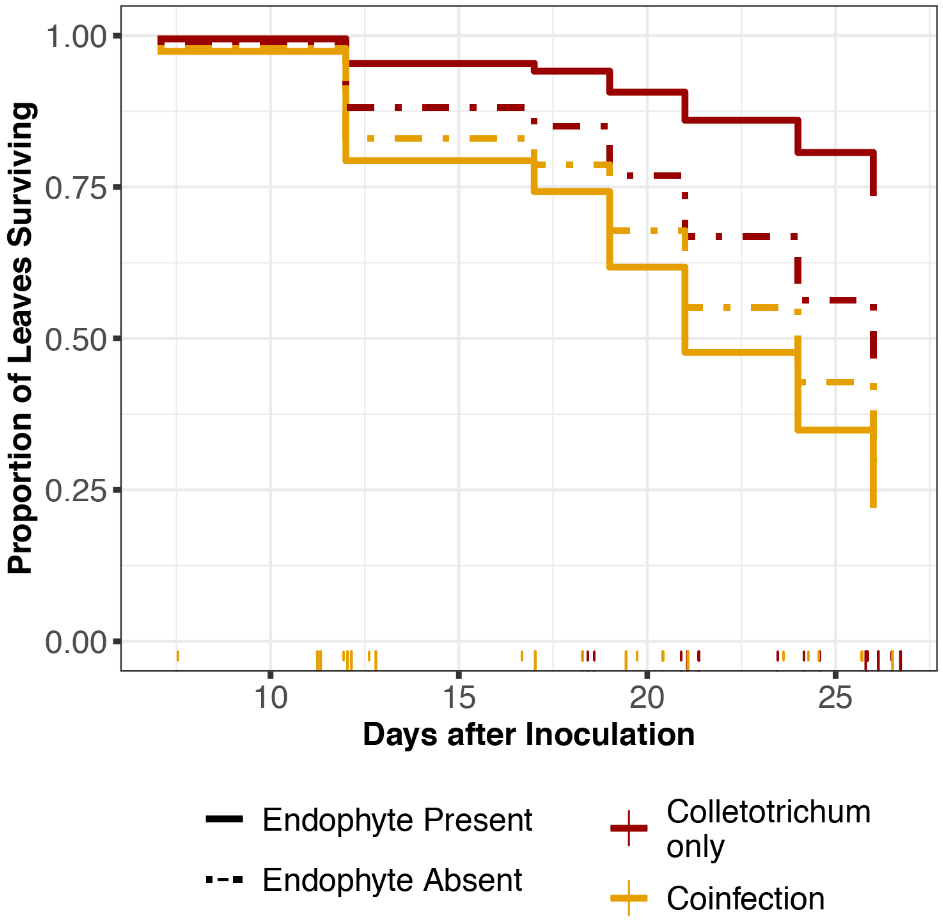


**Figure S2:** Model-estimated relative risk of leaf death over time. Plot is the result of Cox proportional hazards models. For leaves that died, vertical lines along the x-axis show when (in days after inoculation) each leaf died. X-axis values of the vertical lines are jittered to show the data. While there was no difference in leaf survival between leaves inoculated with just *R. solani* and leaves co-inoculated with *R. solani* and *C. cereale* in endophyte-free plants ($p = 0.18$), leaves co-inoculated with both parasites had a 4.92 times higher hazard of mortality than leaves inoculated with only *R. solani* in endophyte-infected plants ($p = 0.008$).

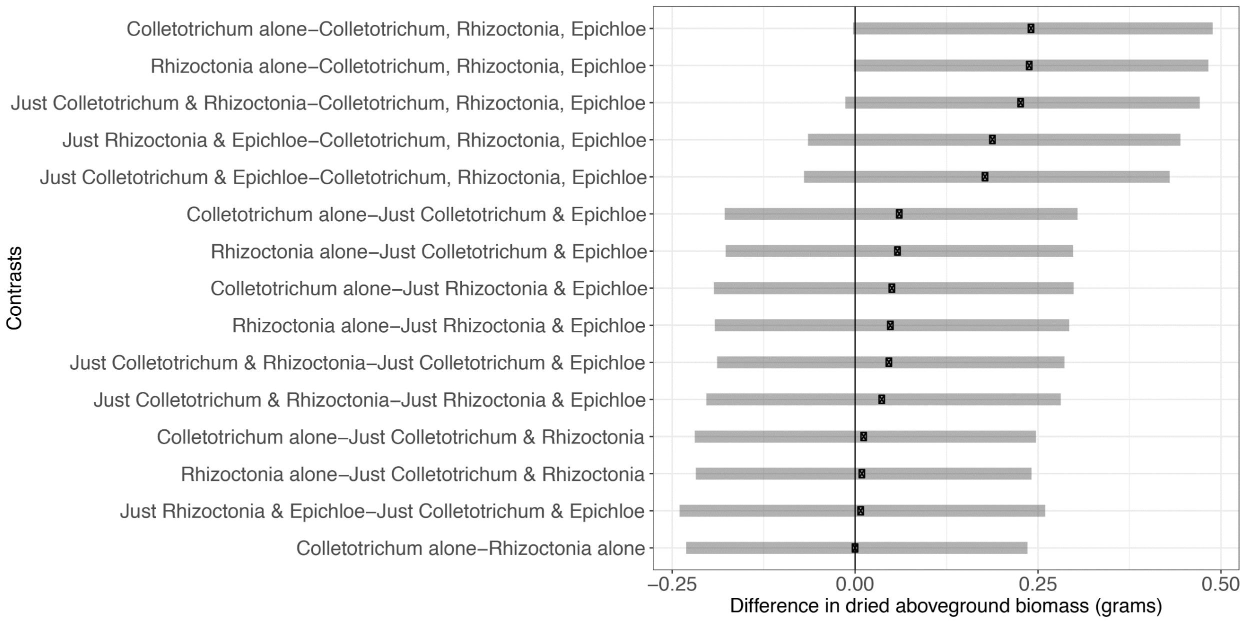


**Figure S3 Plants had lowest biomass when coinfected with *E. coenophiala, C. cereale,* and *R. solani.*** Each row is a contrast as evaluated by the Tukey's Honest Significant Difference method. Points are estimated differences between treatment means with gray bands indicating 95% confidence intervals.

**Table S1: Experiment 1 Setup**

| **Experimental Treatments** | | **Number of plants at start of experiment** | **Number of plants included in analyses (after confirming endophyte presence)** |
|---|---|---|---|
| **Endophyte Treatment** | ***R. solani* treatment** | | |
| Inoculated | Isolate 1 | 19 | 10 |
| | Isolate 2 | 19 | 13 |
| | Isolate 3 | 19 | 14 |
| | Mock | 4 | 0* |
| Free | Isolate 1 | 10 | 10 |
| | Isolate 2 | 10 | 10 |
| | Isolate 3 | 10 | 10 |
| | Mock | 4 | 0* |

*Mock-inoculated control plants were excluded from analyses because no symptoms of parasite infection were observed on them.

**Table S2: Experiment 2 Setup**

| **Parasite Treatment** | **Number of Plants*** | **Sample Size Included in Analyses** |
|---|---|---|
| *C. cereale* alone | 10 | 10 |
| *R. solani* alone | 10 | 10 |
| Simultaneous co-inoculation | 10 | 10 |
| Sequential co-inoculation: *R. solani* first | 10 | 10 |
| Sequential co-inoculation: *C. cereale* first | 10 | 10 |
| Mock-inoculated Control | 10 | 0** |

*Two leaves assayed per plant

**Mock-inoculated control plants were excluded from analyses because no symptoms of parasite infection were observed on them.

**Table S3: Experiment 3 Setup**

| Experimental Treatment | | Number of plants at start of experiment | Number of plants included in analyses (after confirming endophyte presence/lack of symptoms on controls) |
|---|---|---|---|
| **Endophyte Treatment** | **Parasite Treatment** | | |
| Inoculated | *C. cereale* alone | 23 | 14 |
| | *R. solani* alone | 22 | 14 |
| | Coinfection | 22 | 14 |
| | Mock | 4 | 0* |
| Free | *C. cereale* alone | 15 | 15 |
| | *R. solani* alone | 16 | 16 |
| | Coinfection | 16 | 16 |
| | Mock | 4 | 0* |

*Mock-inoculated control plants were excluded from analyses of parasite growth because no symptoms of parasite infection were observed on them. After confirming endophyte presence, 2 endophyte-infected plants and 4 endophyte-free plants were included in analyses of leaf survival.

**Table S4: Experiment 1: *E. coenophiala* vs. *R. solani***

| | ANOVA: Linear mixed effects model of *R. solani* disease progression | | | |
|---|---|---|---|---|
| | **numDF** | **denDF** | **F** | **P** |
| **Intercept** | 1 | 300 | 377.57 | p < 0.0001 |
| **Chamber:Shelf** | 3 | 62 | 0.903 | 0.45 |
| **Days After Inoculation (DAI)** | 1 | 300 | 132.88 | p < 0.0001 |
| **Strain** | 2 | 300 | 27.06 | p < 0.0001 |
| **Endophyte** | 1 | 62 | 2.40 | 0.13 |
| **DAI:Strain** | 2 | 300 | 0.53 | 0.60 |
| **DAI:Endophyte** | 1 | 300 | 0.30 | 0.59 |

**Table S5: Experiment 2: *C. cereale* vs. *R. solani***

**A.**

| | **Linear mixed effects model of *R. solani* disease progression (log transformed)** | | | |
|---|---|---|---|---|
| | **numDF** | **denDF** | **F** | **P** |
| **Intercept** | 1 | 522 | 619.49 | p < 0.0001 |
| **Chamber:Shelf** | 3 | 33 | 0.51 | 0.680 |
| **Days After Inoculation (DAI)** | 1 | 522 | 109.48 | p < 0.0001 |
| **Parasite Treatment** | 3 | 33 | 2.32 | 0.093 |
| **DAI:Parasite Treatment** | 3 | 522 | 3.22 | 0.022 |

**B.**

| | **Post-hoc test of interaction between DAI and inoculation treatment linear mixed effects model of *R. solani* disease progression** | | | |
|---|---|---|---|---|
| **Contrast** | **Estimate** | **SE** | **t** | **P** |
| Rhizoctonia alone – Simultaneous | -0.04 | 0.02 | -2.53 | 0.05 |
| Rhizoctonia alone – Sequential: Rhiz. First | -.02 | 0.02 | -0.85 | 0.83 |
| Rhizoctonia alone – Sequential: Rhiz. Second | -0.05 | 0.02 | -2.51 | 0.05 |
| Simultaneous – Sequential: Rhiz. First | 0.03 | 0.02 | 1.73 | 0.31 |
| Simultaneous – Sequential: Rhiz. Second | -0.003 | 0.02 | -0.20 | 1.00 |
| Sequential: Rhiz. First – Sequential: Rhiz. Second | -0.03 | 0.02 | -1.77 | 0.29 |

**Table S6: Experiment 3 (*Rhizoctonia* results): Factorial Manipulation of *E. coenophiala, C. cereale, and R. solani***

| | **Linear mixed effects model of *R. solani* lesion length (square-root transformed)** | | | |
|---|---|---|---|---|
| | **numDF** | **denDF** | **F** | **P** |
| **Intercept** | 1 | 217 | 544.87 | < 0.0001 |
| **Chamber:Shelf** | 3 | 50 | 1.76 | 0.17 |
| **Days After Inoculation (DAI)** | 1 | 217 | 48.90 | < 0.0001 |
| **Parasite Treatment** | 1 | 50 | 0.56 | 0.46 |
| **Endophyte** | 1 | 50 | 0.43 | 0.51 |
| **Endophyte:Parasite Treatment** | 1 | 217 | 0.51 | 0.51 |
| **DAI:Parasite Treatment** | 1 | 217 | 8.01 | 0.005 |
| **DAI:Endophyte** | 1 | 217 | 0.01 | 0.94 |
| **DAI:Coinfection:Endophyte** | 1 | 217 | 2.57 | 0.11 |

**Table S7: *C. cereale* symptom development**

**A.**

**Survival analysis: *C. cereale* symptoms**

| ***C. cereale Symptom Development Risk (Cox mixed models)*** | **Coefficient** | **Hazard (exponentiated coefficient)** | **Coefficient SE** | **Pr(>\|z\|)** |
|---|---|---|---|---|
| **Reference: *single inoculation with C. cereale*** | | | | |
| Co-inoculation | -0.8599 | 0.4232 | 0.3548 | < 0.001 |
| **Reference: *Endophyte free*** | | | | |
| Endophyte-infected | -0.2078 | 0.8124 | 0.3456 | 0.516 |

**B.**

| ***C. cereale Symptom Development Risk (Cox mixed model)*** | **Coefficient** | **Hazard (exponentiated coefficient)** | **Coefficient SE** | **Pr(>\|z\|)** |
|---|---|---|---|---|
| ***Among leaves from endophyte-infected plants Reference: single inoculation with C. cereale*** | | | | |
| Co-inoculation | -0.8053 | 0.4469 | 0.4705 | 0.0870 |
| ***Among leaves from endophyte-free plants Reference: single inoculation with C. cereale*** | | | | |
| Co-inoculation | -0.9013 | 0.4060 | 0.3442 | 0.0089 |

**Table S8: Survival Analysis Results**

**A.**

| ***Leaf Mortality Risk (Cox mixed model)*** | **Coefficient** | **Hazard (exponentiated coefficient)** | **Coefficient SE** | **Pr(>\|z\|)** |
|---|---|---|---|---|
| ***Reference: control plants*** | | | | |
| *R. solani* alone | 1.64 | 5.17 | 1.03 | 0.03 |
| *C. cereale* alone | 1.17 | 3.22 | 1.04 | 0.08 |
| Co-inoculation | 2.07 | 8.00 | 1.02 | 0.01 |

*Wald statistic=12.34 on 3df, p = 0.006

**B.**

| ***Leaf Mortality Risk (Cox mixed model)*** | **Coefficient** | **Hazard (exponentiated coefficient)** | **Coefficient SE** | **Pr(>\|z\|)** |
|---|---|---|---|---|
| ***Among leaves from endophyte-infected plants Reference: single inoculation with R. solani*** | | | | |
| Co-inoculation | 0.6626 | 1.9397 | 0.4759 | 0.0118 |
| ***Among leaves from endophyte-free plants Reference: single inoculation with R. solani*** | | | | |
| Co-inoculation | 0.1955 | 1.2159 | 0.4372 | 0.581 |

*Wald statistic=8.59 on 3df, p = 0.04

**C.**

| ***Leaf Mortality Risk (Cox mixed model)*** | **Coefficient** | **Hazard (exponentiated coefficient)** | **Coefficient SE** | **Pr(>\|z\|)** |
|---|---|---|---|---|
| ***Among leaves from endophyte-infected plants Reference: single inoculation with C. cereale*** | | | | |
| Co-inoculation | 1.5927 | 4.9172 | 0.5941 | 0.008 |
| ***Among leaves from endophyte-free plants Reference: single inoculation with C. cereale*** | | | | |
| Co-inoculation | 0.3909 | 1.4783 | 0.4607 | 0.1801 |

*Wald statistic=15.48 on 3df, p = 0.001

**Table S9: Survival Analysis Results**

**A.**

| ***Leaf Mortality Risk (Cox mixed model)*** | **Coefficient** | **Hazard (exponentiated coefficient)** | **Coefficient SE** | **Pr(>\|z\|)** |
|---|---|---|---|---|
| ***Reference: control plants*** | | | | |
| *R. solani* alone | 1.49 | 4.43 | 1.02 | 0.0885 |
| *C. cereale* alone | 1.13 | 3.09 | 1.04 | 0.1297 |
| Co-inoculation | 1.52 | 4.58 | 1.04 | 0.0741 |
| **Disease Severity (Percent leaf damaged)** | 0.05 | 1.06 | 0.007 | < 0.001 |

*Wald statistic=115.8 on 4df, p < 0.001

**B.**

| ***Leaf Mortality Risk (Cox mixed model)*** | **Coefficient** | **Hazard (exponentiated coefficient)** | **Coefficient SE** | **Pr(>\|z\|)** |
|---|---|---|---|---|
| ***Among endophyte-infected leaves Reference: single inoculation with R. solani*** | | | | |
| Co-inoculation | 0.23 | 1.26 | 0.51 | 0.62 |
| ***Among endophyte-free leaves Reference: single inoculation with R. solani*** | | | | |
| Co-inoculation | -0.27 | 0.75 | 0.46 | 0.43 |
| **Disease Severity (Percent leaf damaged)** | 0.05 | 1.05 | 0.01 | < 0.001 |

*Wald statistic=138.7 on 4df, p < 0.001

C.

| ***Leaf Mortality Risk (Cox mixed model)*** | **Coefficient** | **Hazard (exponentiated coefficient)** | **Coefficient SE** | **Pr(>\|z\|)** |
|---|---|---|---|---|
| ***Among endophyte-infected leaves Reference: single inoculation with C. cereale*** | | | | |
| Co-inoculation | 1.21 | 3.35 | 0.62 | 0.006 |
| ***Among endophyte-free leaves Reference: single inoculation with C. cereale*** | | | | |
| Co-inoculation | -0.32 | 0.72 | 0.48 | 0.48 |
| **Disease Severity (Percent leaf damaged)** | 0.05 | 1.06 | 0.01 | < 0.001 |

*Wald statistic=90.89 on 4df, p < 0.001

**Table S10: Experiment 3 Analysis of aboveground biomass of plants**

| | Biomass | | | |
|---|---|---|---|---|
| | numDF | denDF | F | P |
| **Chamber:Shelf** | 3 | 92 | 13.17 | < 0.001 |
| **Parasite Treatment** | 2 | 93 | 1.58 | 0.22 |
| **Endophyte Treatment** | 1 | 94 | 5.69 | 0.02 |
| **Parasite Treatment:Endophyte Treatment** | 2 | 93 | 1.63 | 0.20 |

**Table S11: Experiment 3: Biomass means and standard errors**

| Parasite/Endophyte Treatment | Mean Biomass (g) | SE |
|---|---|---|
| *Colletotrichum* alone | 0.791 | 0.058 |
| *Rhizoctonia* alone | 0.807 | 0.056 |
| Just *Colletotrichum* and *Rhizoctonia* | 0.811 | 0.056 |
| Just *Rhizoctonia* and *Epichloë* | 0.757 | 0.062 |
| Just *Colletotrichum* and *Epichloë* | 0.802 | 0.061 |
| *Colletotrichum*, *Rhizoctonia*, and *Epichloë* | 0.666 | 0.062 |
